# Newborns’ sensitivity to speed changes as a building block for animacy perception

**DOI:** 10.1101/2020.11.17.386466

**Authors:** Elisa Di Giorgio, Marco Lunghi, Giorgio Vallortigara, Francesca Simion

## Abstract

The human visual system can discriminate between animate beings *vs*. inanimate objects on the basis of some kinematic cues, such as starting from rest and speed changes by self-propulsion. The ontogenetic origin of such capability is still under debate. Here we investigate for the first time whether newborns manifest an attentional bias toward objects that abruptly change their speed along a trajectory as contrasted with objects that move at a constant speed. To this end, we systematically manipulated the motion speed of two objects. An object that moves with a constant speed was contrasted with an object that suddenly increases (*Experiment 1*) or with one that suddenly decreases its speed (*Experiment 2*). When presented with a single speed change, newborns did not show any visual preference. However, newborns preferred an object that abruptly increases and then decreases its speed (*Experiment 3*), but they did not show any visual preference for the reverse sequence pattern (*Experiment 4*). Overall, results are discussed in line with the hypothesis of the existence of attentional biases in newborns that trigger their attention towards some visual cues of motion that characterized animate perception in adults.

## Introduction

Motion of living things–i.e., *animate motion* – is a rich source of cues about their behaviors, goals and intentions, and humans, as well as other animals, attend to these cues [1,2,3,4]. Animacy perception, a perceptual phenomenon which reflects visual processing that is specialized for the extraction of animacy from visual cues of motion [5, 6], can be studied with simple stimuli showing the motion of just one or two objects [7, 8, 9, 10, 11, 12, 13, 14]. Heider and Simmel’s [15] classic study is the most compelling evidence of such a phenomenon, demonstrating that a “fairly fast, automatic, irresistible and highly stimulus-driven” sense of animacy could arise from basic dynamic perceptual cues. Among the visual cues of motion that trigger animacy perception in adults, the most powerful are self-propulsion (e.g., an entity’s capacity to initiate motion without an applied external force), motion contingency, discontinuity in motion trajectory, the violation of Newtonian laws, body-axis alignment, and sudden direction and speed changes without any contact with other objects [8, 9, 12, 13, 16].

From a developmental perspective, the irresistible and automatic nature of animacy perception in adults based on these simple low-level perceptual visual cues of motion raises questions on its ontogenetic origins. Whether the animate-inanimate distinction is present early in life, what perceptual cues signal to the infant that an entity is animate and what is the role of visual experience, are relevant and intriguing open questions. The animate-inanimate distinction is well in place in the first year of life. Broadly speaking, infants understand very early that animate entities behave in ways in which inanimate objects do not [17, 18, 19, 20, 21, 22, 23, 24, 25, 26]. Very young infants not only categorize animate entities on the basis of start from rest and speed changes by self-propulsion [26, 27, 28], but also attribute goals and intentions to objects that move on their own [29, 30, 31, 32, 33]. Further, chasing perception in 4- and 10-month-old infants is mainly triggered by the presence of sudden accelerations and by the specific relations between the movements of the two objects [34].

However, controversy still surrounds the question about the developmental origin of animacy perception. Some authors credit the infant with innately based, domain-specific systems that are sensitive to certain perceptual visual cues of motion (i.e., self-propulsion) in identifying agents and animate beings [20, 22, 35, 36]. For instance, Mandler has hypothesized that motion provides infants with conceptual knowledge about the “kinds of things” objects are [36]. In the same way, other authors have hypothesized that at birth, infants possess separate core knowledge systems for animate and inanimate objects, and that they understand that constraints on motion that apply to physical objects may not hold true for animate beings [22, 37, 38]. As an alternative to this domain-specific view, other authors presented a theoretical framework that has at its core the idea that general rather than specific mechanisms are sufficient to account for how infants perceive and process the motion properties that characterized an object as animate or inanimate (i.e., the *constrained attentional associative learning framework*, 39, 40, 41). Specifically, according to this more perceptually oriented view, infants use learning mechanisms that operate across all domains of knowledge to encode statistical regularities in their environment: they would thus learn about properties of the objects that appear in a self-propelled event through their experiences with specific kinds of entities that engage in these motions. A third theoretical framework, the *cue-based bootstrapping model*, proposed that animacy perception develops by bootstrapping from an inborn sensitivity to general low-level visual cues of motion, into a learning mode central to infants’ later concept of animacy that appears later during development. In this view, the presence of inborn predispositions to some visual cues of motion that characterized animate beings might be a sort of jumpstart to the development of animacy perception through learning mechanisms [42].

The aim of the present paper was to investigate whether a sensitivity to some rudimentary visual cues of motion that in older infants and adults elicited animacy perception is present at birth or is the result of learning and experience-dependent processes.

Differently from adult studies where kinematic cues are manipulated together to elicit the highest level of animacy perception [13], here the main aim was to investigate the role of each of those individual visual cues of motion in attracting the newborns’ attention, put them under separate experimental control. Using such method, recent comparative studies in newborns and chicks demonstrated that sensitivity to some cues of motion that trigger animacy perception in adults are present very early in life. For example, visually self-propelled objects whose motion reveals an internal energy source of energy attract naïve chicks [43]. Chicks also are attracted by moving objects that can accelerate and decelerate by their own [44]. They were presented with two animated-event stimuli, in each one a red ball appeared on the screen already in motion (i.e., no cues about the onset motion were available), moved linearly along the horizontal trajectory and then disappeared at the opposite side of the screen. In the “speed-change stimulus”, the object changes its speed two times: it suddenly accelerated at one third of the way and then suddenly decelerated to the original speed at two thirds of the way. This stimulus was compared to the “constant-speed stimulus”, where the object moved at a constant speed. Results showed that naïve chicks show a preference to an object that changes its speed over an object that moves at a constant speed. Intriguingly, the preference disappeared when the exact moment of the sudden speed change is occluded.

As for newborns, it has been demonstrated that they are sensitive to certain well-defined perceptual spatio-temporal cues present in a physical causal event (i.e., launching event) [45]. Recently, Di Giorgio and colleagues showed for the first time that newborns are sensitive to a start from rest by self-propulsion [46]. Newborns were presented with two-computer generated events in which the sudden onset motion of an object without an applied external force, and therefore perceived as self-propelled by adults, was manipulated. In that study, an event in which an object suddenly started from rest without the application of external forces was clearly visible to the baby, was contrasted with an ambiguous event, in which the object appeared already on the screen in motion so as no cues about the onset motion were available. In that scenario, newborns showed a spontaneous visual preference for the object that abruptly started from rest by self-propulsion, demonstrating an early sensitivity to this visual cue of motion that triggers animacy perception in adults [46].

At present, it is not known whether human newborns are sensitive to other elementary motion cues that elicit animacy perception in adults, besides start from rest without any external force. Indeed, objects that change their velocity without an apparent external cause, tend to be perceived as animate and intentional. On the other hand, objects that do not change velocity or change their motion on the base of external causes are judged as inanimate. Moreover, objects perceived as moving relatively faster are perceived as animate more often than relatively slower objects [12], and a sudden deceleration tends to favour the perception of animacy less than a corresponding sudden acceleration [13].

Due to speed change relevance as a visual cue of motion in eliciting animacy perception in adults [7, 8, 13], here we sought to investigate whether a sensitivity to an object that changes its speed on its own, without any interaction with an external object, is present at birth.

To test this hypothesis, newborns were presented simultaneously with two computer-generated events in which we systematically manipulated only one visual cue of motion that is known to elicit animacy perception in adults, that is the speed of an object, in four different visual preference experiments. In Experiment 1 and 2, a single speed change was tested. Experiment 1 comprised an object that after a third of its own trajectory suddenly increases its velocity and continues its trajectory with such motion pattern speed. In Experiment 2, an object that after a third of its own trajectory suddenly diminished its velocity and continued its trajectory with such motion pattern speed was presented. Then, we tested newborns’ sensitivity to two consecutive speed changes, Experiment 3 comprised an object that at one third of its trajectory suddenly increases and after another third diminished its velocity, returning to its original speed. Experiment 4, comprised an object that at one third of its trajectory suddenly diminished and after another third increased its velocity, returning to its original speed. The control stimulus was always an object that moves at a constant speed (that we called “speed-constant stimulus”) (Figure 1).

The dependent variables recorded were the mean fixation duration towards the stimuli and the visual preference percentage towards the speed-change stimulus. In addition, as in a previous study from our lab [46], we analysed the location where newborns were looking at the exact time when the object changes its speed (for details see Methods and Supplementary Information).

**Figure 1:**
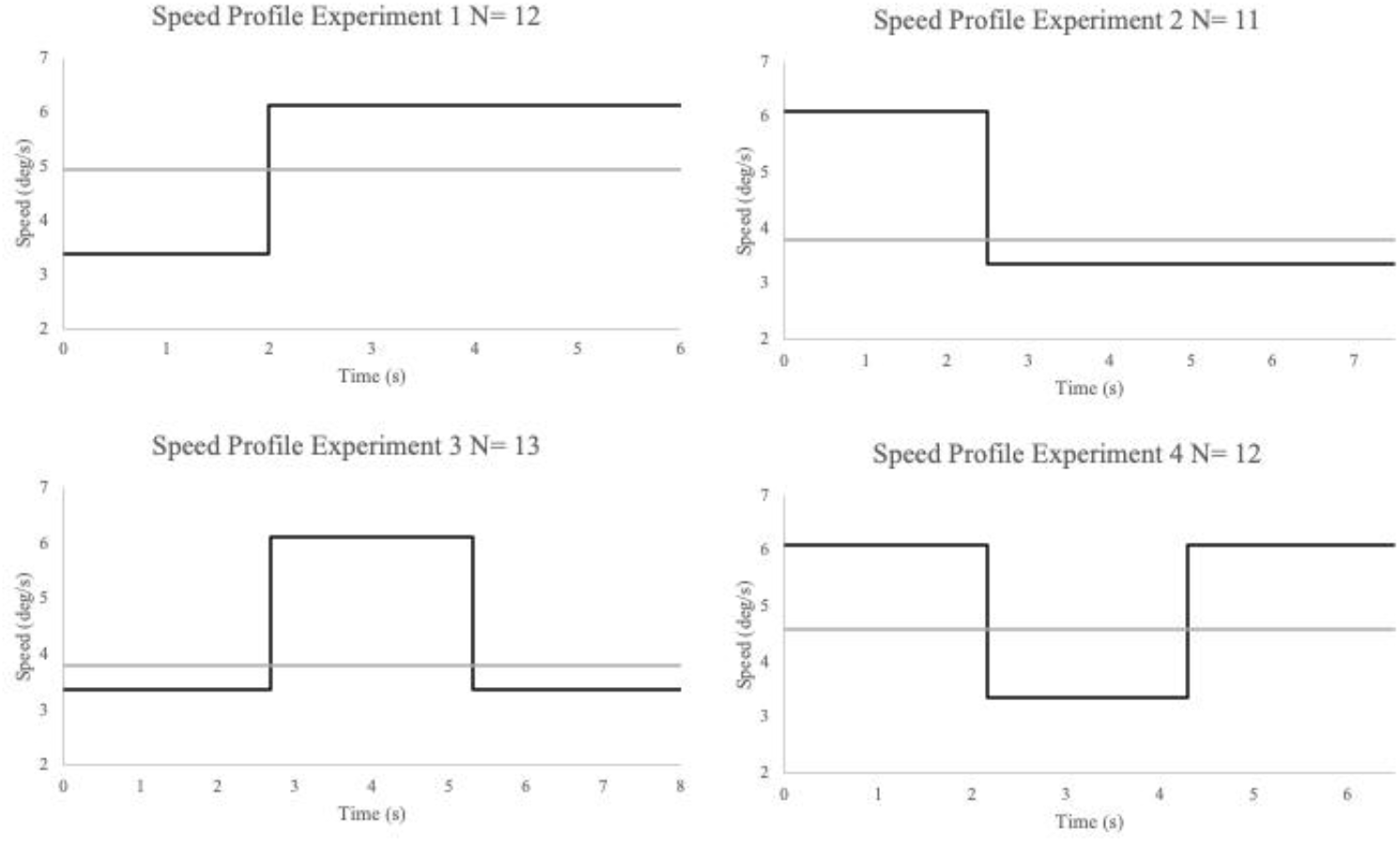
Schematic representation of stimuli employed in the four Experiments. In each graph, the velocity of the object in the speed-changes event *vs*. the velocity of the object in the constant speed event as a function of time was plotted. The black line represents the speed change stimuli, whereas the gray line represents the constant speed stimuli.

## Results

Results showed that when stimuli comprised a single speed-change *(Experiment 1 and 2*), they did not trigger newborns’ attention. Specifically, newborns did not look longer at the stimulus that abruptly increases (*M_mean duration fixation_* = 3.73 s, *SD* = 1.36 *vs*. “speed-constant stimulus”, *M_mean duration fixation_* 4.95 s, *SD* = 3.62, *t*_11_ = 1.43, *p* = 0.18, n.s.) or decreases its speed (*M_mean duration fixation_* 3.94 s, *SD* = 1.50 *vs*. “speed-constant stimulus”, *M_mean duration fixation_* = 4.13 s, *SD* = 1.83, *t*_10_=0.54, *p* = 0.67, n.s.). In line with this, newborns did not show any visual preference for the speed-change stimulus neither in Experiment 1 (*M_Percentage score_* = 47.80%, *SD* = 10.82, *t*_11_ = 0.72, *p* = 0.49, n.s.) nor in Experiment 2 (*M_Percentage score_* = 49.4%, *SD* = 11.30, *t*_10_= 0.19, *p* = 0.86, n.s.). Seven out of 12 newborns preferred to look at the increased speed stimulus (binomial test, *p* = 0.77, n.s.), and only 4 out of 11 newborns preferred to look at the decreased speed stimulus compared to the speed-constant event (binomial test, *p* = 0.55, n.s.). Further, the analyses on the location where newborns were looking at the exact time when the object changes its speed suggested that they were not able to detect a single and abrupt speed change, since the percentage was not above the chance level (*Experiment 1*, *M* = 46.10%, *SD* = 9.10, one sample t-test, *t*_11_ = 1.33, *p* = 0.21) (*Experiment 2*, *M* = 47.31%, *SD* = 7.42, one sample t-test, *t*_10_= 1.26, *p* = 0.23).

Results of Experiment 1 and 2 seem to suggest that a single speed change in terms of increase and decrease speed respectively is not sufficient for eliciting a spontaneous newborns’ visual preference. At a first glance, this suggested that speed changes as perceptual visual cues are not relevant for triggering newborns’ attention such as self-propulsion did [46]. However, even if the null results should be interpreted with caution, they are in line with those studies that demonstrated that only one single speed-change was not a very strong perceptual visual cue in triggering animacy perception in adults [8, 13]. In contrast, it has been demonstrated that two consecutive speed changes (a sudden speed increase followed by an abrupt speed decreased), if they are allocated in coherent and coordinated sequence, were particularly effective in eliciting perception of animacy in adults [8] as well as in chicks [44]. Therefore, the specific aim of Experiments 2 and 3 was to test whether this is the case also for human newborns.

Results suggested that newborns are sensitive to two consecutive speed changes, but only in the presence of an abrupt velocity increase followed by a sudden velocity decrease (*M_mean duration fixation_* = 3.47 s, *SD* = 1.53 *vs*. speed-constant stimulus *M_mean duration fixation_* = 2.43 s, *SD* = 0.57, *t*_12_= 2.38, *p* = 0.035, Cohen’s *d* = 0.66) (*Experiment 3*). Eleven out of the 13 newborns preferred to look at the two consecutive increase-decrease speed change stimulus (binomial test, *p* = .022), and their visual preference for that stimulus was significantly above the chance level, (*M* = 55.10%, *SD* = 7), *t*_12_ = 2.40, *p* = 0.033, Cohen’s *d* = 0.66). In addition, the analyses on the location where newborns were looking at the exact time when the object changes its speed revealed that the percentage was above the chance level (*Condition 3*, *M* = 62.20%, *SD* = 15.10, one sample t-test, *t*_12_ = 2.84, *p* = 0.015, Cohen’s *d* = 0.79). On the contrary, the stimulus with the reverse speed change sequence (abrupt decrease followed by a sudden increase in velocity) did not grab newborns’ attention (*Experiment 4*).

They did not look longer at that stimulus (*M_mean duration fixation_* = 3.64 s, *SD* = 1.07 *vs*. speed-constant stimulus *M_mean duration fixation_* = 3.53 s, *SD* = 1.10, *t*_11_= 0.38, *p* = 0.71, Cohen’s *d* = 0.11). Only 4 out of the 12 newborns preferred to look at the two consecutive decrease-increase speed change stimulus (binomial test, *p* = .038), and their visual preference for that stimulus was not significantly above the chance level, (*M* = 49.32%, *SD* = 12.20, one sample t-test, *t*_11_ = 0.32, *p* = 0.89). Finally, the analyses on the location where newborns were looking at the exact time when the object changes its speed revealed that the percentage was not above the chance level (*M* = 51.30%, *SD* = 16.0, one sample t-test, *t*_11_ = 0.16, *p* = 0.875) (Figures 2, 3 and 4).

**Figure 2:**
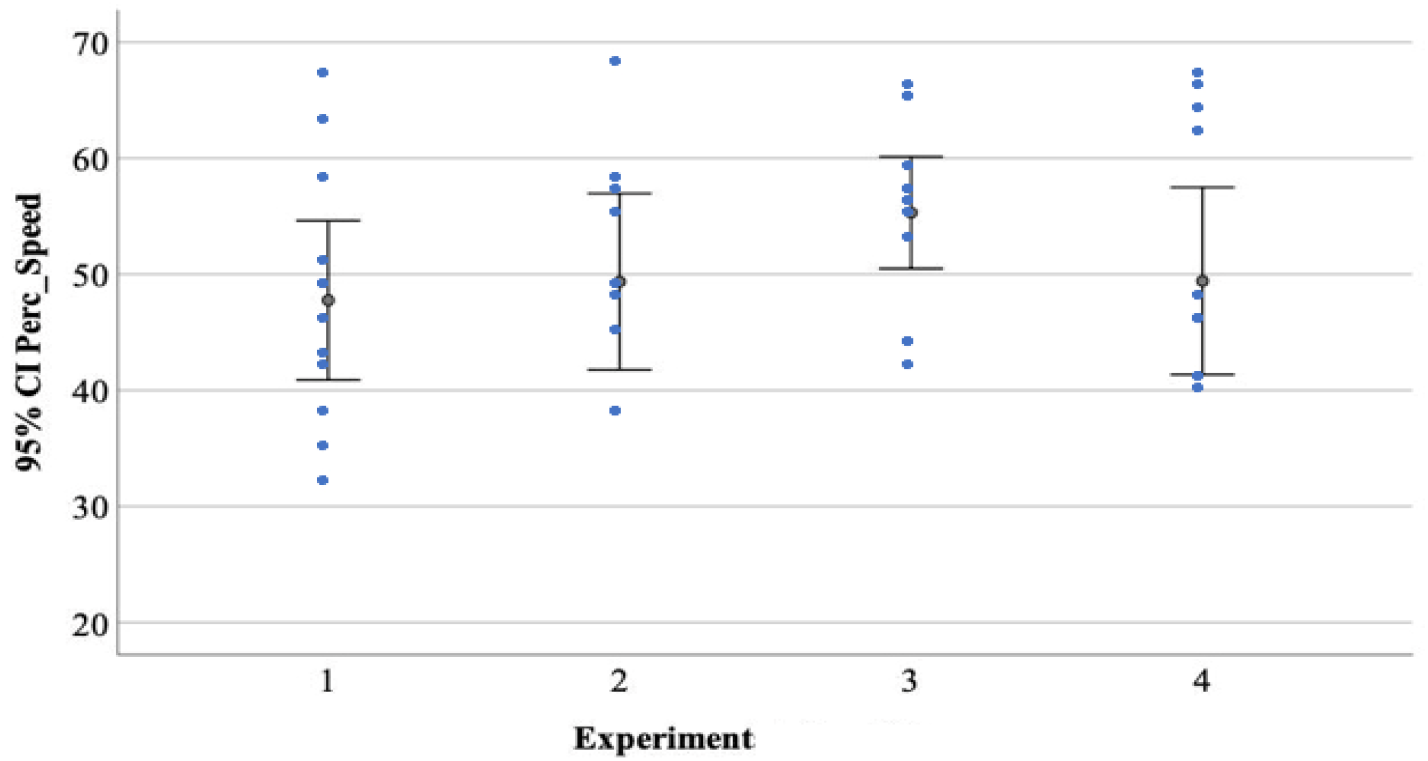
Average percentage of visual preference for the speed-change event in the four Experiments. The error bars represent the 95% Confidence Interval for the mean value.

**Figure 3:**
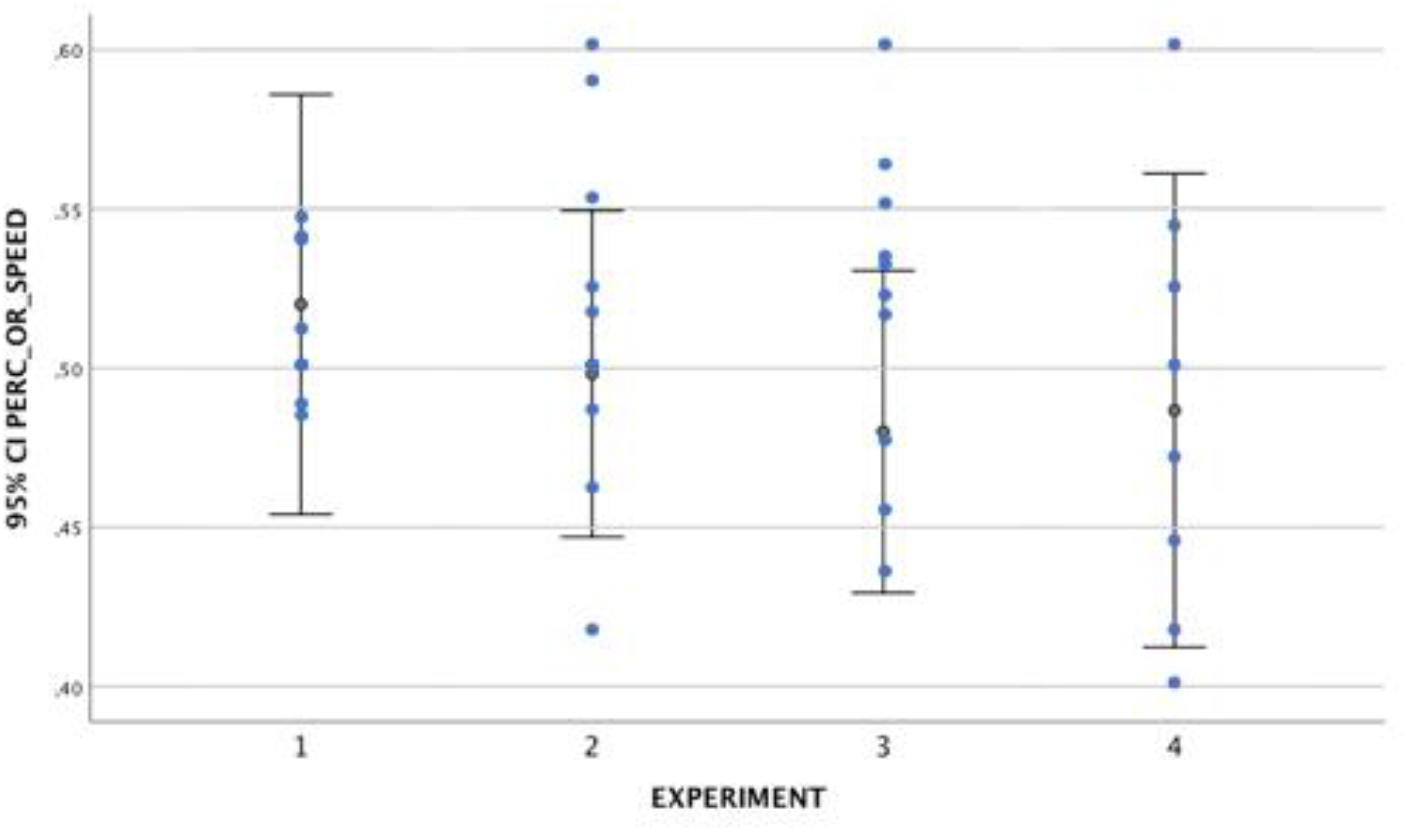
Average percentage of number of orienting responses for the speed-change event in the four Experiments. The error bars represent the 95% Confidence Interval for the mean value.

**Figure 4:**
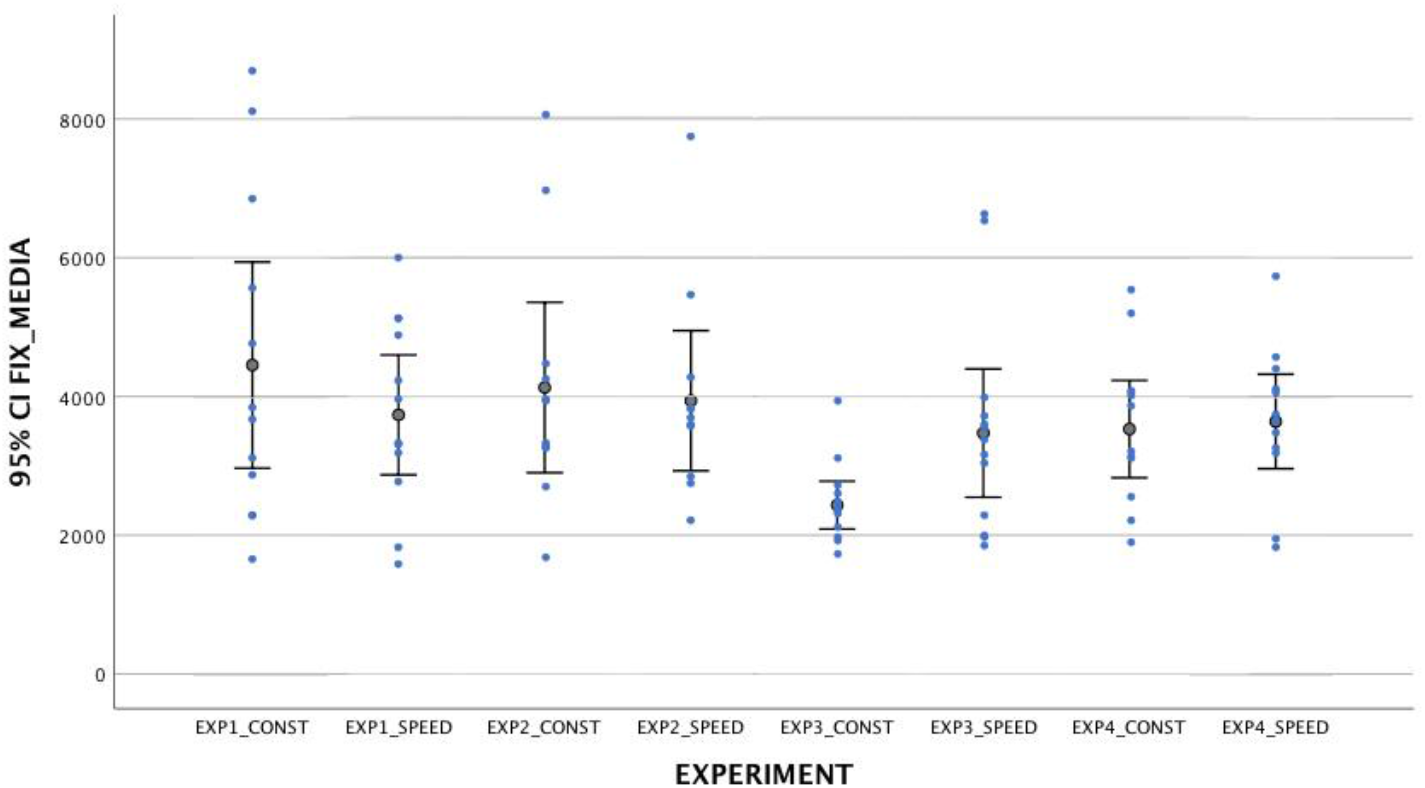
Mean duration fixation toward speed-change and constant speed stimuli in the four Experiments. The error bars represent the 95% Confidence Interval for the mean value.

Overall, results seemed to suggest that newborns have some rudimentary attentional bias toward speed-changes, but they need to have sufficient information, since they preferred the object that changed its speed twice.

## Discussion

Inferring animacy is the developmental precursor to complex social understanding. While there is mounting evidence for the precocious acquisition of the animacy concept, much less is known about perceptual visual cues infants use to acquire such an “abstract” concept from early on. Among dynamic cues, start from rest by self-propulsion is critical to trigger animacy perception in adults [13], and recent comparative studies demonstrated that sensitivity to this visual motion cue is present from birth in chicks [43] and human newborns [46]. In addition to such visual cues of motion, in the present work we demonstrated that human newborns’ attention is triggered also by stimuli that abruptly change their speed without any contact with external forces or objects.

In line with those studies that demonstrated that the presence of one change in speed does not favor much the perception of animacy in adults [8,13], we found that a single change in speed is not sufficient to elicit a visual preference in newborns (Experiment 1 and 2). On the contrary, newborns looked longer at the object that suddenly increased and then decreased its speed compared to the same object that moves at a constant speed (*Experiment 3*). Further, when the object abruptly decreased and then increased its speed (Experiment 4), newborns did not show any visual preference. Overall, results suggest that newborns have some rudimentary attentional bias toward speed-changes, but they need to have sufficient information (e.g. two consecutive abrupt speed changes), and that the ordering of the speed change has a role in eliciting their visual preference. Apparently, an abrupt speed increase, followed by a sudden speed decrease, is a powerful combination of speed changes for eliciting newborns’ preference.

An interesting point concerns the fact that, in order to investigate newborns’ sensitivity to speed-changes, the stimuli employed here resemble a step acceleration or deceleration change. Indeed, our “speed-change stimulus” abruptly increased or decreased their speed, without any pause. Therefore, to test whether the visual perceptual system at birth is sensitive to acceleration, future studies should implement stimuli, such those employed in the present study, with a gradual speed change, that is the more natural speed profile expected for animate creatures. Indeed, we already know that such stimulus manipulation elicited a spontaneous preference for the natural speed changing stimulus in visually naïve chicks [44], but not in human newborns [47, 48]. However, it must be pointed out that in the study on newborns, the stimuli employed were complex Point-Light Display (PLD) of goal-directed grasping hand actions and, therefore, no direct comparison between the results of the previous and present study can be done.

Overall, the present work showed that newborns respond also to changes in speed of already moving stimuli, in addition to start from rest by self-propelled movement found in a previous study [46], supporting the hypothesis that since birth our species has some attentional biases toward visual cues of motion that trigger animacy perception in adults.

These biases might be considered as the building blocks on which, during the development, infants and adults enhance their ability to discriminate animate from inanimate beings. The present result, if considered together with similar data found in visually naïve chicks [44], demonstrated the presence of attentional predispositions tuned toward one of the general properties of animate creatures that older infants and adults perceived as animate.

Although speed changes and start from rest by self-propulsion are motion cues that elicit newborns’ visual preferences, how these cues interact to promote such preferences is still largely unknown. Therefore, future studies should investigate whether one of these visual cues is more powerful compared to the other, or whether they have additive effects in triggering newborns’ attention.

To conclude, our data are compatible with the idea that an initial sensitivity to low-level visual cues leads infants from birth to attend to visual cues of motion that characterized animate beings [49], and to learn rich correlations among these cues as they parse animates from inanimates (i.e., *cue-based bootstrapping model*, 42). This learning appears to be central to infants’ later concept of animacy, guiding their socio-emotional understanding, the theory of mind, and predictions of action [17, 26, 40]. The extent to which experience is needed for the expression of animacy distinction should be tested in future.

Understanding how our visual system perceives animacy is a mandatory step in understanding how we perceive the social world. The perception of animacy is an automatic process that begins in early infancy, but in some cases this perception develops atypically, leading to difficulties in perceiving animacy, such as in the case of autism spectrum disorders [50, 51] (see some recent studies on an animal model of autism [52]). Therefore, knowing more about the quality of motions that triggers such a fundamental perceptual ability is vital to understanding how our social cognition works in both typical and atypical individuals.

## Methods

### Participants

A total of 61 healthy and full-term newborns were tested (31 males). Thirteen babies were excluded from analyses because they showed a position bias (n = 5) (i.e., he looked in one direction more than 80% of the total fixation time) [46, 53] or because they became tired and started crying during testing (n= 8).

Therefore, the final sample comprised 48 (28 males) healthy and full-term newborns. They were randomly assigned to one of the four Experiments: n = 12 in *Experiment 1*, n =11 in *Experiment 2*, n = 13 in *Experiment 3*, and n = 12 in *Experiment 4*. Their postnatal age ranged from 12 to 139 h (*M_age_* = 43.4 h, *SD* = 27.8). All subjects met the normal delivery screening criteria, had a birth weight between 2440 g and 4105 g (*M* = 3325 g, SD =440.5), and had an Apgar score > 9 at 5 min. Newborns were tested only if awake and in an alert state, and after the parents had provided informed consent. All the experiments described have been carried out in accordance with the Declaration of Helsinki. Further, the Azienda Ospedale - Università degli Studi di Padova has licensed all experimental procedures (Protocol number: 91644645).

### Stimuli

Stimuli consisted of two video-animation events displayed on an Apple LED Cinema Display (Flat Panel 30”) computer monitor (refresh rate = 60 Hz, 2560 × 1600), representing the movement of an object (i.e., a gray ball of 3 cm of diameter already used in previous studies with human newborns, [45, 46]. The baby sat on an experimenter’s lap at a distance of approximately 30 cm from the computer screen, therefore at such distance 1 degree of visual angle is equivalent to 0.53 cm. In each of the four Experiments, two stimuli were presented simultaneously on the screen, on the right and one on the left. In both events, the object always entered the newborn’s view already in motion, appearing from the external limit of a white background, therefore no cues about “start from rest” were available. Similarly, once completed its motion, the object disappeared from view while still moving. After the entrance, the object always moved along a straight trajectory towards the center of the screen. One stimulus was always characterized by the presence of visible speed-changes (i.e., speed increment or decrement, and both) during his trajectory, whereas the other moved at a constant speed (see Figure 1). Videos were produced by looping the animations.

#### Experiment 1

An object that increases its speed without the contact of external forces and a speed-constant stimulus were presented. In the speed-change stimulus, the object suddenly increased its speed at one-third (5.3 cm, 10.02°) of the distance (16 cm, 28.07°). The speed changed from 1.76 cm/s to 3.2 cm/s. In contrast, in the speed-constant stimulus the object covered the same distance (16 cm) at a constant speed of 2.6 cm/s. Both events described lasted 6 s (150 frames, 25 frames/s).

#### Experiment 2

The speed-change stimulus comprised an object that suddenly decreases its speed without any contact with external forces at one-third (5.3 cm, 10.02°) of the distance (16 cm, 28.07°). The speed changed from 3.2 cm/s to 1.76 cm/s. In contrast, in the speed-constant stimulus, the object covered the same distance (16 cm) at a constant speed of 2 cm/s. Both events described lasted 7.5 s (188 frames, 25 frames/s) (see Figure 1 and 5).

**Figure 5:**
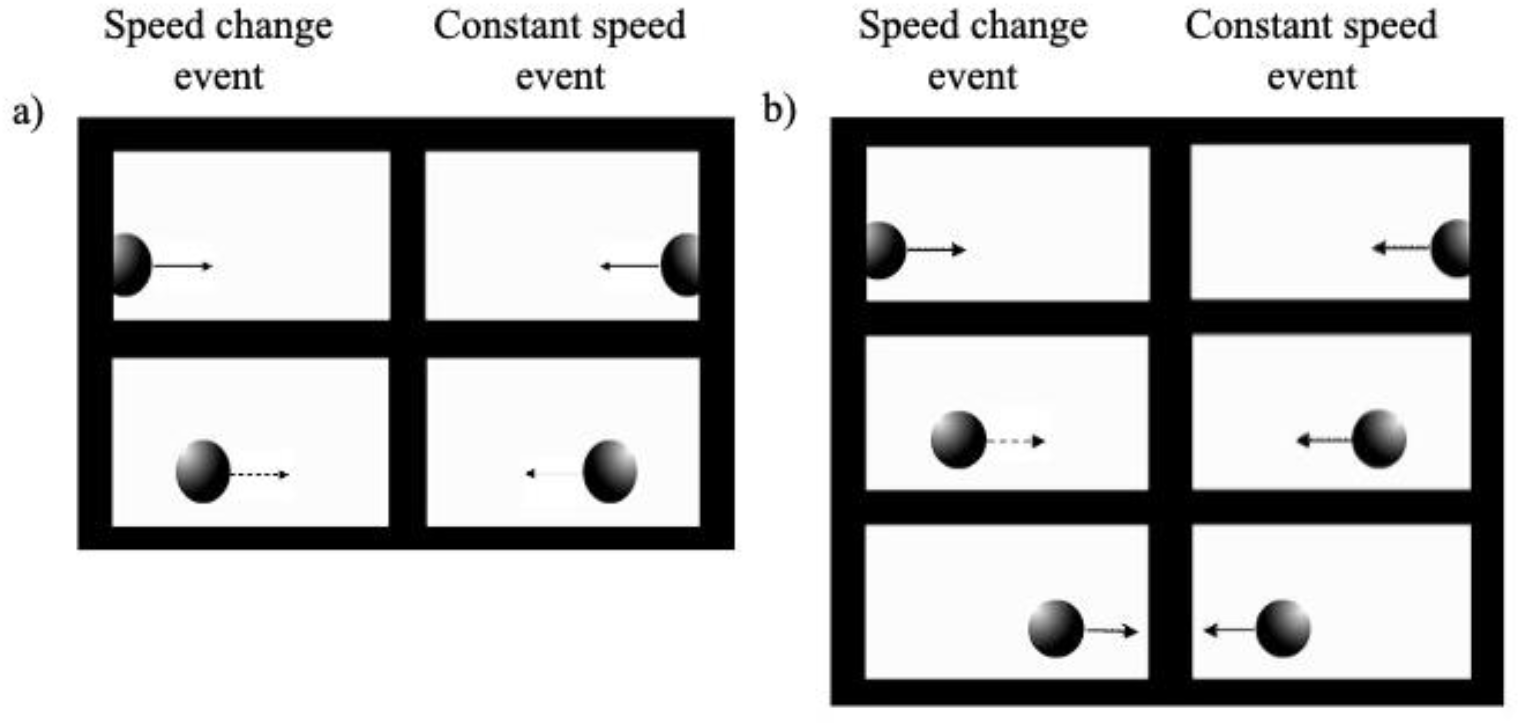
Schematic representation of the events used in the four Experiments. Panel a) Experiment 1 and 2, left sequence represents the Speed-Change Event and right sequence the Constant Speed Event. Panel b) Experiments 2 and 3, left sequence represents the Speed-Change Event and right sequence the Constant Speed Event.

#### Experiment 3

The speed change stimulus was an object that suddenly increased its speed at one third (5.3 cm, 10.02°) of the total distance (16 cm, 28.07°), and then decreased it back to the original speed at two third of the way (10.6 cm, 19.46°). The speed changed from 1.76 cm/s to 3.2 cm/s and then returned at 1.76 cm/s. In contrast, in the constant-speed motion event, the object covered the same distance (16 cm) at a constant speed of 2 cm/s. Both events described lasted 8 s (200 frames, 25 frames/s).

#### Experiment 4

An object that abruptly decreased its speed at one third (5.3 cm, 10.02°) of the total distance (16 cm, 28.07°) and then increased it back to the original speed at two third of the way (10.6 cm, 19.46°), characterized the speed-change stimulus. In this case, the speed changed from 3.2 cm/s to 1.76 cm/s and then returned at 3.2 cm/s. In contrast, in the constant-speed motion event, the object covered the same distance (16 cm) at a constant speed of 2.4 cm/s. Both events described lasted 6.5 s (163 frames, 25 frames/s) (see Figure 1 and 5).

To note, the low speed values adopted here derive from previous studies. For instance, it has been shown [54], using a visual preference procedure, estimated thresholds at about 1.4deg/s at 4 weeks of age, and 6.7 deg/s at 10 weeks [55,56].

### Apparatus and Procedure

We employed the infant-control preferential looking technique [57]. Stimuli presentation and data collection were performed using E-Prime 2.0. The baby sat on an experimenter’s lap in front of the screen and white curtains were drawn on both sides of the newborn to prevent interference from irrelevant distractors. Above the monitor, the video camera recorded the eye movements of newborns to control their looking behavior on-line and to allow off-line coding of their fixations [46] (see Figure 6).

**Figure 6:**
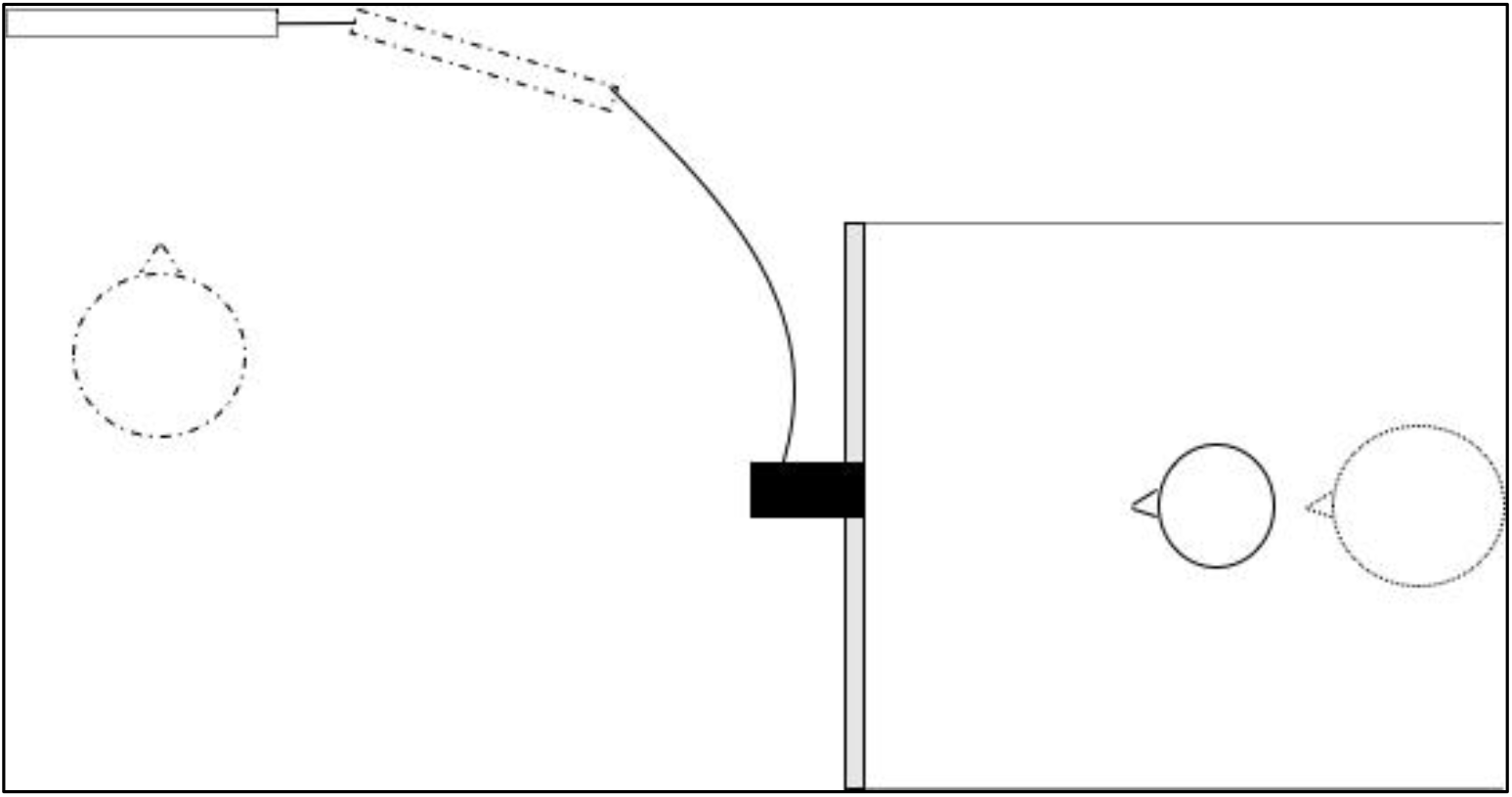
Schematic representation of the Lab Set up during data collection. On the right, the black line circle represents the baby sitting on the experimenter’s lap (the dot line circle) at a distance of 30 cm from the computer screen (the thin grey rectangle). At such a distance 1 degree of visual angle is equivalent to 0.53 cm. Above the monitor, the video camera (the black square connected to a monitor behind the newborns’ computer screen) recorded the eye movements of newborns. On the left, the second experimenter (the dashed line circle) manages the stimuli presentation with E-prime 2.0 installed on a second computer (black line rectangle).

At the beginning of each experimental condition, a red disc on a black background appeared to attract the newborn’s gaze to the centre of the monitor (i.e., fixation point). In a continuous fashion, the disc changed in size from small (diameter of 1.8 cm) to large (diameter of 2.5 cm), and vice-versa until the newborn’s gaze was properly aligned with the red disc. The red disk blinked at a rate of 300 ms on, and 300 ms off. The sequence of trials was then started by a second experimenter who watched the newborn’s eyes through the monitor. When the newborn’ s gaze was aligned with the red disk, the second experimenter pressed a keyboard key that automatically turned off the central disc activated the onset of the stimuli and a timer on the video, thereby initiating the sequence of trials. Because stimuli were presented bilaterally on the left and the right side of the monitor with a convergent motion pattern (from peripheral to the central visual field), each newborn was given two paired presentations (trials 1 and 2) of the test stimuli. In each trial, the position of the stimuli was reversed (the initial left-right order of presentation was counterbalanced across subjects). A trial ended when the newborn did not fixate on the display for at least 10 s. When the newborn shifted his/her gaze away from the stimuli for more than 10 sec, the stimuli were removed and the disc was turned on to start Trial 2. The experiment finished when the newborn shifted his/her gaze away from the stimuli for more than 10 sec during Trial 2.

The four Experiments were planned and conducted as completely independent experiments, and thus, no direct comparison among the experiments was applicable and no correction for the alpha level in any of the statistical tests was conducted for each of the experiments.

We recorded both the baby’s eye-movement by a camera in front of the stimuli presented and the trial sequence in order to permit the coding off-line frame by frame. Further, a timer on the video that starts exactly at the same time when the first trial starts allows us to synchronize the video of the baby’s eyes and the stimuli’s sequence to connect the exact moment on the frame to the presented stimuli.

Two coders, unaware of the stimuli presented, analyzed off-line the videos by coding newborns’ eye movements frame-by-frame. The average level of inter-observer agreement was Pearson’s *r* = 0.93, *p* < 0.001.

According to Cohen’s model of infants’ attention [58], the dependent variables that we recorded are i) the total number of orienting responses towards the stimuli as index of the attention getting mechanisms. This index is calculated as the sum of how many times newborns orient their gaze towards either the stimulus on the left or the stimulus on the right in both trials, and ii) the mean duration of fixation towards the stimuli as an index of the attention holding mechanisms. Moreover, we calculated the percentage of visual preference in each experimental condition, defined as the total fixation time (TFT) toward the speed-change stimulus divided by the total fixation time spent in looking at both stimuli (TFT speed-change / TFT speed-change + TFT constant-speed) and compared this percentage to the chance level (50%). Scores significantly above 50% indicate a visual preference for the speed-change stimulus. These analyses are traditionally performed with data from newborns [50, 59] To note, results on the total number of orienting responses missed statistical significance and therefore we did not report them (but see Supplementary Information).

In addition, we decided to conduct further analyses in order to investigate whether the sudden speed changes motion per se attracted newborns’ attention in each experimental condition. We analysed the location where newborns were looking at the exact time when the object changes its speed. To do so, we calculated the total number of orienting responses towards the object that changes its speed and divided this number by the total number of orienting responses towards both stimuli at the same exact time, X 100 [46]. Since in Experiment 3 and 4 there were two consecutive speed changes each and newborns who looked at the first change remained on that stimulus and looked also at the second one (*Experiment 3*, *t*_12_ = 0.26, *p* = 0.80 and *Experiment 4*, *t*_11_= 0.90, *p* = 0.39), we collapsed together the two speed changes.

### Data availability

The dataset of the present study is available from the corresponding author on reasonable request.

## Acknowledgements

The authors are deeply indebted to Professor Eugenio Baraldi, Director of the Intensive Care Unit and Professor Giorgio Perilongo Director of the Department of Women and Child Health for their collaboration. Special thanks to the nursing staff of the Pediatric Clinic (Azienda Ospedaliera) to the babies who took part in the study and to their parents, to Elena Berto, Daniela Carà, and Anna Bianchi for assistance with newborn testing. This study was carried out within the framework of the agreement between the University and the Azienda Ospedaliera of Padova prot.: 91644645. The research was supported from the European Research Council under the European Union’s Seventh Framework Programme (FP7/2007-2013) Grant ERC-2011-ADG_20110406, Project No: 461 295517, PREMESOR to G.V. and supported by Fondazione Caritro Grant Biomarker DSA [40102839] to G.V., PRIN 2018, Project No: 20152M5A5J, to G.V.

## Author contributions statement

E.D.G., M. L., G.V. and F.S. designed the study; M. L. designed the stimuli; E.D.G. and M.L. collected and analyzed the data, and did the statistical analyses; E.D.G interpreted the data and wrote the manuscript; M. L., G.V. and F.S. discussed the results and critically revised the manuscript.

## Competing interests

The authors declare no competing financial interests.

## Notes

### Competing Interest Statement

The authors have declared no competing interest.

