## Supplementary Information for "Newborns’ sensitivity to speed changes as a building block for animacy perception"

**Additional statistical analyses**

We conducted additional analyses on the total number of orienting responses that indexed attention-getting mechanisms [1]. Here we reported the results for each experimental condition.

*Experiment 1:*

Increased speed stimulus ($M$ = 17.7, $SD$ = 5.7) *vs.* constant-speed stimulus ($M$ = 17.1, $SD$ = 6.9), $t_{11}$ = 0.30, $p$ = 0.77.

*Experiment 2:*

Decreased-speed stimulus ($M$ = 13.4, $SD$ = 4.2) *vs.* constant-speed stimulus ($M$ = 13.8, $SD$ = 5.3), $t_{10}$ = 0.33, $p$ = 0.75.

*Experiment 3*:

Stimulus that increased and the decreased its speed ($M$ = 15.7, $SD$ = 6.8) *vs.* constant-speed stimulus ($M$ = 16.3 $SD$ = 5.3),

$t_{12}$ = 0.44, $p$ = 0.67.

*Experiment 4:*

Stimulus that decreased and then increased its speed ($M$ = 13.3, $SD$ = 4.9) *vs.* constant-speed stimulus ($M$ = 13.8, $SD$ = 4.6),

$t_{11}$ = 0.26, $p$ = 0.80.
